# FL/FLT3 signaling enhances mechanical pain hypersensitivity through Interleukin-1 beta (IL-1β) in male mice

**DOI:** 10.1101/2025.04.09.648037

**Authors:** Corinne Sonrier, Chamroen Sar, Ivo Melo, Lucie Diouloufet, Gabriel V. Lucena-Silva, Sylvie Mallié, Juliette Bertin, Jean-Philippe Leyris, Fabrice Ango, Thiago M Cunha, Hamid Moha ou Maati, Jean Valmier, Cyril Rivat, Ilana Méchaly

**Affiliations:** Université de Montpellier, Montpellier, France; Inserm U-1298, Institut des Neurosciences de Montpellier, Montpellier, France; Center for Research in Inflammatory Diseases (CRID), Department of Pharmacology, Ribeirão Preto Medical School, University of Sao Paulo (USP), Ribeirão Preto, Brazil; IMmune HEAlth International Laboratory (IMHEA, IRL, CNRS), Ribeirão Preto, Brazil

**Keywords:** pain, neuroinflammation, cytokine, sensory neurons, tyrosine kinase receptor

## Abstract

Fms-like tyrosine kinase 3 (FLT3) plays a critical role in chronic pain through its ligand FL, a cytokine that triggers mechanical pain hypersensitivity. However, the underlying molecular mechanisms remain unclear. Here, we investigate the potential interplay between FL and IL-1β a key cytokine in DRG neurons sensitization and mechanical hyperalgesia through both *in vitro* and *in vivo* approaches. ELISA assays reveal that intrathecal FL administration significantly increases IL-1β protein levels in both the DRG and dorsal spinal cord of mice, beginning four hours post-injection. Using video microscopy and [Ca^2+^]_i_ fluorescence imaging in primary DRG neuron cultures, we demonstrate that FL potentiation of TRPV1 receptor responses to capsaicin is partially mediated by IL-1β signalling, as evidenced by a significant reduction in this potentiation in the presence of the IL-1 receptor antagonist, IL-1Ra. Furthermore, FLT3-driven acute mechanical pain hypersensitivity *in vivo* is reduced both by prior administration of IL-1Ra and in IL-1 receptor knockout mice. Importantly, IL-1β-induced mechanical pain hypersensitivity remains independent of FLT3 signalling as shown in *Flt3* knockout mice. Collectively our findings expand the understanding of neuro-immune interactions by demonstrating a potential functional link between FL/FLT3 and IL-1β/IL-1R signalling in nociceptive processing.

**Highlights:** Cytokines are known to engage in complex interactions and regulate each other

FL increases IL-1β in DRG and DSC within 4h, but IL-1β does not affect FL levels

FL modulates capsaicin-induced Ca^2+^ influx *via* IL-1β/IL-1R signalling in DRG neurons

IL-1R inhibition delays or abolishes FL-induced mechanical hypersensitivity *in vivo*

**Graphical abstract:** The timeline of the experimental design and the graphical abstract were created with BioRender.com

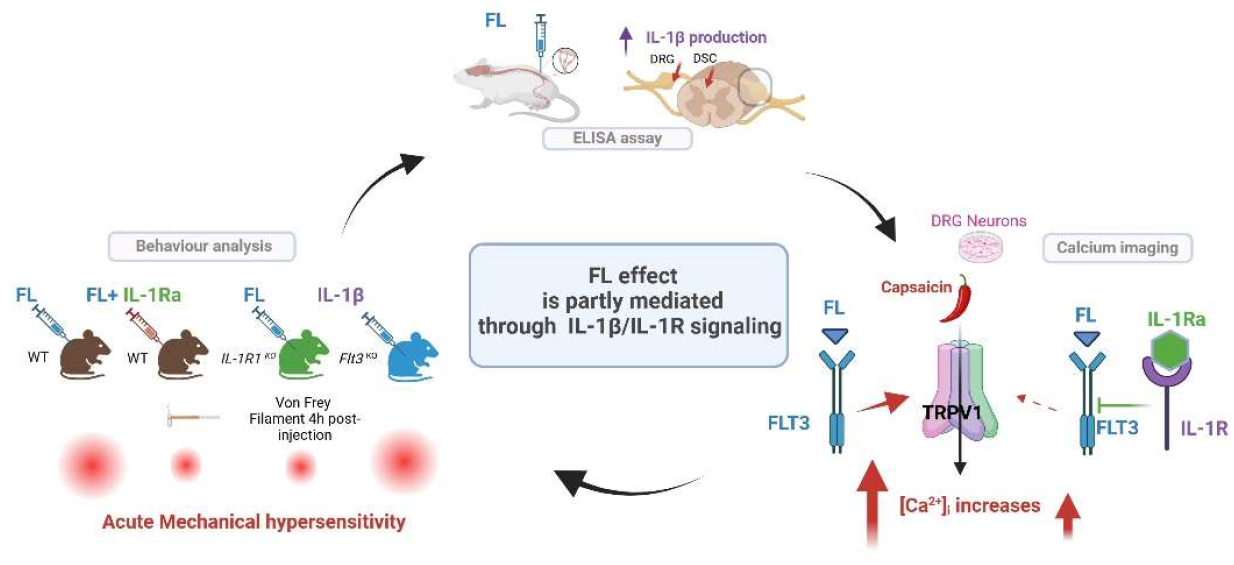

## 1. Introduction

The intricate interactions between the nervous and immune systems play a critical role in both health and disease. Bidirectional communication between these systems is essential during development for maintaining homeostasis, responding to environmental changes, and regulating stress and disease (Dantzer, 2018; Chu et al., 2020).

Chronic pain is recognized as a neuro-immune disorder characterized by elevated levels of proinflammatory mediators. The complex interactions between immune cells and neurons within the dorsal root ganglion (DRG) and spinal cord are key regulators of neuronal sensitization, leading to long-term hyperexcitability promoting persistent pain (Marchand et al., 2005; Moalem and Tracey, 2006; Grace et al., 2010).

While the cytokine FL, Fms-like tyrosine kinase 3 (FLT3) Ligand, and its cognate receptor FLT3 have been extensively studied for their role in haematopoiesis, our investigations have highlighted the critical role of FLT3 receptor expressed in DRG sensory neurons in the development and persistence of peripheral neuropathic pain (PNP) (Rivat et al., 2018). Targeting FLT3 specifically in DRG sensory neurons has been demonstrated to replicate or reverse various symptoms associated with nerve injury. Notably, this includes the potentiation of TRPV1 channels by FL in cultured DRG neurons and cellular and molecular long-term *in vivo* modifications in DRG neurons, resulting in hyperexcitability and PNP-like symptoms. Conversely, specific genetic or pharmacological inhibition of FLT3 in DRG sensory neurons has been shown to alleviate PNP symptoms (Rivat et al., 2018).

Furthermore, FL immunoreactivity was found to be colocalized with CD45, a marker of haematopoietic cells, at the nerve injury site in mouse. This suggests the presence of infiltrating CD45-positive cells expressing FL following PNP (Rivat et al., 2018). Despite shedding light on FLT3 role in PNP, the exact cellular and molecular mechanisms by which FLT3-expressing sensory neurons contribute to the sensitization of pain pathways remain largely unexplored.

The overproduction of the proinflammatory cytokine interleukin-1β (IL-1β) is a central event involved in the pathophysiological changes observed in various disease states, including chronic pain. IL-1β exerts its effects primarily *via* the interleukin-1 receptor type 1 (IL-1R1), which mediates intracellular signalling cascades. In contrast, interleukin-1 receptor type 2 (IL-1R2) functions as a decoy receptor, binding IL-1β without initiating signal transduction (McMahan et al., 1991; Orlando et al., 1997). IL-1β whose role in the development of mechanical hyperalgesia is multifaced may increase the expression of other pro-nociceptive factors such as Nerve Growth Factor, initiate signalling pathways that promote the release or activation of pain-related molecules and/or directly influence neuronal excitability through interaction with ion channel receptors including TRPV1 or NMDA (Ren and Torres, 2009). In addition, IL-1R1 has been shown to be expressed by sensory neurons, especially in a subpopulation of TRPV1 positive nociceptors and its activation promotes increased excitability of DRG nociceptors (Binshtok et al., 2008; Mailhot et al., 2020).

Considering the key contribution of FL/FLT3 and IL-1β/IL-1R signalling in the sensitization of DRG sensory neurons, the present study explores their potential interaction using both *in vitro*, and *in vivo* investigations.

## 2. Material and Methods

### Ethics statement

All the procedures carried out in this study were approved by the French Ministry of Research (authorization #1006).

### Animals

Experiments were performed on C57BL/6 naive mice (Janvier, France) or mice carrying a homozygous deletion of *Flt3* (*Flt3*^*KO*^; Mackarehtschian et al., 1995) or *Il1r1* (*Il1r1*^*KO*^ mice; The Jackson Laboratory, Glaccum et al., 1997) and their respective littermates (*Flt3*^*WT*^ and *Il1r1*^*WT*^) weighing 25–30 g. Animals were maintained in a climate-controlled room on a 12-hour light/dark cycle and had ad libitum access to food and water. Housing conditions were kept at a temperature of 22–23°C and a humidity level of 40–50%

### Behavioural analysis

Before testing, mice were acclimatized for 60 min in the temperature and light-controlled testing room within a plastic cylinder or on wire mesh. Experimenters were blinded to the genotype, or the drug administered. Tactile withdrawal threshold was determined as previously described (Rivat et al., 2018). A 0.6 g-von Frey filament was used to test hind-paw mechanical hypersensitivity. Sharp withdrawal of the stimulated hind-paw was considered as a positive response. The procedure was applied 10 times and the percentage of positive responses was calculated.

### Adult sensory neuron culture

For calcium imaging, neuron cultures were established from lumbar (L4–L6) DRG (Elzière et al., 2014). Briefly, ganglia were successively treated by two incubations with collagenase A (1 mg/ml, Roche Diagnostic, France) for 45 min each (37 °C) and trypsin-EDTA (0.25%, Sigma, St Quentin Fallavier, France) for 30 min. They were mechanically dissociated through the tip of a fire-polished Pasteur pipette in neurobasal (Life Technologies, Cergy-Pontoise, France) culture medium supplemented with 10% fetal bovine serum and DNase (50 U/ml, Sigma). Isolated cells were collected by centrifugation and suspended in neurobasal culture medium supplemented with 2% B27 (Life Technologies), 2 mM glutamine, penicillin/streptomycin (20 U/ml, 0.2 mg/ml) plated at a density of 2500 neurons per coverslip and were incubated in a humidified 95% air 5% CO_2_ atmosphere at 37 °C. The data were obtained from experiments conducted on at least two biological replicates.

### Calcium imaging

For calcium imaging video microscopy [Ca^2+^] _i_ fluorescence imaging, DRG neurons were loaded with fluorescent dye 2.5 µM Fura-2 AM (Invitrogen, Carlsbad, CA) for 30 min at 37 °C in standard external solution contained: 145 mM NaCl, 5 mM KCl, 2 mM CaCl_2_, 2 mM MgCl_2_, 10 mM HEPES, 10 mM glucose (pH adjusted to 7.4 with NaOH and osmolarity between 300 and 310 mOsm). The coverslips were placed on a stage of Zeiss Axiovert 200 inverted microscope (Zeiss, München). Observations were made at room temperature (20–23 °C) with a ×20 UApo/340 objective. Fluorescence intensity at 505 nm with excitation at 340 and 380 nm were captured as digital images (sampling rates of 0.1–2 s). Regions of interest were identified within the soma from which quantitative measurements were made by re-analysis of stored image sequences using MetaFluor Ratio Imaging software. [Ca^2+^]_i_ was determined by ratiometric method of Fura-2 fluorescence from calibration of series of buffered Ca^2+^ standards. Neurons were distinguished from non-neuronal cells by applying 25 mM KCl, which induced a rapid increase of [Ca^2+^]_i_ only in neurons. Pulses of capsaicin were applied at 2 min-intervals and capsaicin and drugs was added after the third pulse. The same temporal stimulation protocol was used for TRPV1 (capsaicin). All drugs and solutions were applied with a gravity-driven perfusion system. For data analysis, amplitudes of [Ca^2+^]_i_ increases, Δ*F*/*F*_max_, caused by stimulation of neurons with capsaicin were measured by subtracting the “baseline” *F*/*F*_max_ (mean for 30 s prior to capsaicin addition) from the peak F/Fmax achieved on exposure to capsaicin. In the absence of any treatment the distribution of these ratios was well fitted by a normal distribution.

### Production of human recombinant FL (rh-FLT3-L)

Recombinant FL was produced in the *E. coli* Rosetta (DE3) strain (Novagen) in our laboratory using the pET15b-rhFL plasmid according to the protocol described (Verstraete et al., 2009) with some minor modifications. The rh-FL was checked for endotoxin content using the Pyrogen Recombinant Factor C endotoxin detection assay from LONZA (Walkersville MD, USA) and was found free of endotoxins.

### Drugs delivery

FL, IL-1β, IL-1Ra were intrathecally injected (Rivat et al., 2018). Briefly, a 30 G needle attached to a microsyringe was inserted between L4 and L5 vertebrae in lightly restrained, unanaesthetized mice. The reflexive tail flick was used to confirm the punction. A total volume of 10 μl was injected.

### IL-1β and FL cytokines Elisa based assay measurement

After collection of DRG and dorsal horn of the spinal cord (DSC) from naïve animals, tissues were snap-frozen in liquid nitrogen and stored at −80°C until processing for ELISA. DRG and DSC were homogenized in the MagNA Lyser Instrument (Roche; twice for the speed of 6000 for 30 s) with ceramic spheres (Lysing Matrix D, MP Biomedicals, #116913100) in lysis buffer (10 mM Tris-HCl, pH 7.4, 50 mM NaCl, 1 % Triton X100, 1 mM EDTA, 1 mM EGTA, DTT 1 mM) supplemented with complete protease inhibitor cocktail (Sigma-Aldrich #P8340). Then, centrifugation (10 000 rpm for 15 min at 4°C) was performed. Supernatants were collected. Mouse IL-1β/IL-1F2 Quantikine ELISA kit (RnD Systems, Minneapolis, United States, MLB00C) and Mouse/Rat Flt3 Ligand Quantikine ELISA Kit MFK00 were used for the measurement of total IL-1β and FL amounts in DRG and DSC according to manufacturer recommendations. Data were calculated per mg of proteins.

### Statistical analyses

All experiments were randomized. Data are expressed as the mean ± s.e.m. All sample sizes were chosen based on our previous studies except for animal studies for which sample size has been estimated via a power analysis using the G-power software. The power of all target values was 80% with an alpha level of 0.05 to detect a difference of 50%. Statistical significance was determined by analysis of variance (ANOVA one-way or two-way for repeated measures, over time). In all experiments in which a significant result was obtained, the F test was followed by Bonferroni post-hoc test for multiple comparisons. In case of two experimental groups, unpaired two-tailed t-test was applied. For ELISA quantification, statistical analyses were performed using Mann–Whitney test. The applied statistical tests are specified in each figure legend.

## 3 RESULTS

### 3.1 FL induces overexpression of IL-1β in mice lumbar DRG and dorsal spinal cord

To investigate the potential interaction between FL and IL-1β in the nociceptive pathways, we conducted ELISA assays to measure the levels of both cytokines in lumbar DRG and DSC following intrathecal injections of either cytokine or saline four hours post-injection. Results, illustrated in **Figure 1**, reveal that recombinant FL significantly increases IL-1β total protein levels in both the DRG and DSC from four hours post-injection compared to the saline-injected groups (**Figure 1A, B**). Specifically, the basal levels of IL-1β cytokine originally 69.23 ± 12.04 pg/mg in DRG and 19.37 ± 6.11 pg/mg in DSC extracts significantly increased (p<0.05 and p<0.01) to 145.9 ± 19.68 pg/mg in lumbar DRG and 65.23 ± 13.32 pg/mg in the spinal cord extracts, respectively. Conversely, no significant change in FL protein levels was observed following intrathecal IL-1β injection compared to saline-injected controls in both DRG and DSC (**Figure 1C, D**) at the same time post injection. These data indicate that FL induces a transient production of the pro-inflammatory cytokine IL-1β that may contribute to the cellular effects of FL in the nociceptive pathways.

**Figure 1:**
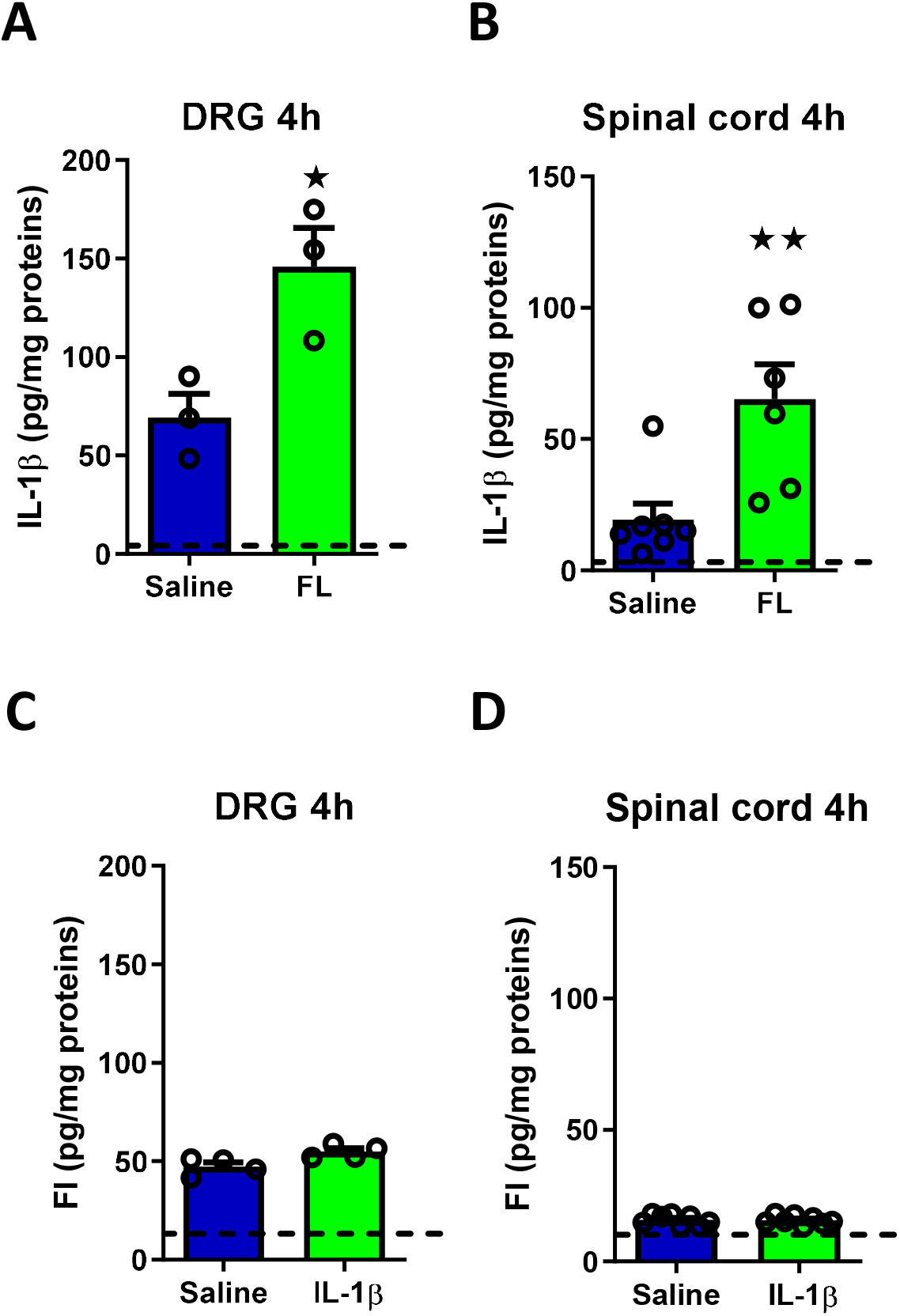
Histograms showing ELISA measurements of IL-1β or FL levels after intrathecal injection of FL or IL-1β, respectively, at 4 hours in the DRG and spinal cord. **A)** ELISA measurements of IL-1β levels in the DRG 4 hours after intrathecal injection of 13.8 nM FL versus saline solution. **B)** ELISA measurements of IL-1β levels in the spinal cord 4 hours after intrathecal injection of 13.8 nM FL versus saline solution. **C)** ELISA measurements of FL in the DRG 4 hours after intrathecal of 5 pg/µl of IL-1β injection versus saline solution. **D)** ELISA measurements of FL levels in spinal cord 4 hours after intrathecal of 5 pg/µl of IL-1β injection versus saline solution. Dash line corresponds to the detection limit dosage: 5 pg for IL-1β and 10 pg for FL according to the information provided with the ELISA kit. Number of experiments n = 3 to 8 animals per group, Statistical analysis: Mann–Whitney test ⋆ p value < 0.05; ⋆⋆ p value < 0.01.

### 3.2 FL potentiates TRPV1 function *in vitro* through IL-1β Receptor activation

The TRPV1 receptor plays a crucial role in neuropathic pain, acting as a key mediator in the nociceptive pathway. Its activation is critical for the development and sensitization processes that underlie neuropathic pain (Caterina et al., 2000; Davis et al., 2000; Basbaum et al., 2009). We previously showed that FL alone had no effect on basal [Ca^2+^] _i_, but markedly potentiated, in a concentration-dependent manner, the TRPV1 response to repeated applications of capsaicin, a specific TRPV1 agonist (Rivat et al., 2018).

To examine the functional impact of IL-1β on TRPV1 channel response through IL-1 receptor activation *in vitro*, we investigated the effect of recombinant FL, specifically focusing on the potentiation of calcium (Ca^2+^) influx induced by capsaicin, with or without the presence of the receptor antagonist IL-1Ra. IL-1Ra is a natural inhibitor of IL-1β’s pro-inflammatory effects (Arend, 1991).

Results of video microscopy [Ca^2+^] _i_ fluorescence imaging on primary cultures of mice adult DRG neurons are presented in **Figure 2**. Controls experiments designed to ensure the specificity of IL-1Ra effect on IL-1β activity are presented in **Figure 2 A**. Data show the potentiation of intracellular calcium levels [Ca^2+^] _i_ induced by capsaicin both alone (0.5 µM) or in combination with IL-1β (1pg/µl) in the presence or absence of IL-1Ra (**Fig 2A**). DRG neurons treatment with the receptor antagonist IL-1Ra (0.333 pg/µl) had no effect on [Ca^2+^] _i_ levels induced by capsaicin (0.5 µM) alone and completely inhibited the potentiation of [Ca^2+^] _i_ levels induced by IL-1β (**Fig. 2A**). **Figure 2B** shows that FL (5.6 nM) significantly enhances the capsaicin induced Ca^2+^ response. Our data further show that FL potentiation of [Ca^2+^] _i_ levels induced by capsaicin was significantly reduced in the presence of the antagonist IL-1Ra.

**Figure 2:**
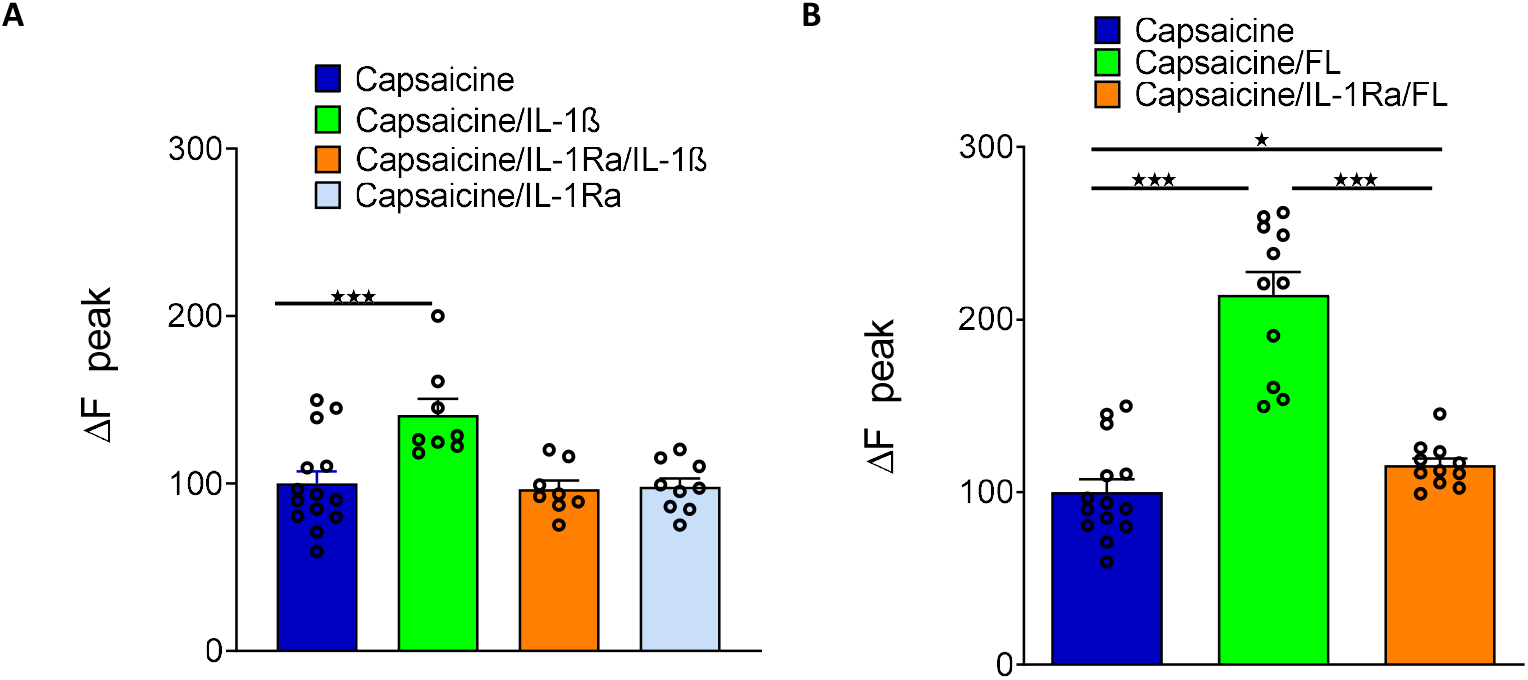
Histograms showing intracellular calcium dynamic in different conditions on primary culture of neurons from DRG using fura-2-am cell calcium imagine technique. **A)** F340/F380 fluorescence ratio calcium response after Capsaicin 0.5 µM (control condition, n=14 cells), Capsaicin 0.5 µM + IL-1β 1 pg/µl (n=8 cells), Capsaicin 0.5 µM + IL-1β 1pg/µl + IL-1Ra 0.333 pg/µl (n=8 cells) and Capsaicin 0.5 µM + IL-1Ra 0.333 pg /µl (n=9 cells) cell perfusion. **B)** F340/F380 fluorescence ratio calcium response after Capsaicin 0.5 µM (control condition, n=14 cells), Capsaicin 0.5 µM + FL 5.6 nM (n=11 cells) and Capsaicin 0.5 µM + FL 5.6 nM + IL-1Ra 0.333 pg/µl (n=11 cells) cell perfusion. Number of experiments n = 2 replicates of primary culture of DRG neurones, statistical analysis: one-way ANOVA followed by Bonferroni test for multiple comparisons, ⋆ p value < 0.05; ⋆⋆⋆ p value < 0.001.

These *in vitro* functional assays indicate that FL effect on capsaicin-induced Ca^2+^ influx is partly mediated through the IL-1β/IL-1R signalling pathway. These findings highlight a potentially significant interaction between FL and IL-1β in the modulation of TRPV1-mediated signalling.

### 3.3 Blocking IL-1R signalling affects FL induced mechanical pain hypersensitivity *in vivo*

Building on previous *in vitro* results indicating a link between FLT3 signalling and the IL-1β/IL-1R pathway, we sought to determine whether the pronociceptive effects of FLT3 activation involve IL-1β signalling *in vivo*. To address this point, we examined the impact of IL-1β signalling blockade on FL*-*induced mechanical pain hypersensitivity *in vivo* using the receptor antagonist IL-1Ra or *Il1r1* knockout mice.

A time-course experiment was conducted to illustrate IL-1β-induced mechanical pain hypersensitivity in the presence or absence of IL-1Ra in mice. Paw withdrawal thresholds were measured over a 6-day period. IL-1β (5pg/10µl) was administered by IT injection one hour after IL-1Ra (1ng/10µl) as shown in the timeline in **Figure 3A**. The graph shows that IL-1β injection significantly increased mechanical pain sensitivity peaking at 48 hours post-injection and returning towards baseline within 6 Days (**Figure 3B)**. The administration of IL-1Ra significantly reduced IL-1β-induced mechanical pain hypersensitivity, confirming the antagonistic action of IL-1Ra on IL-1-induced IL-1R activation.

**Figure 3:**
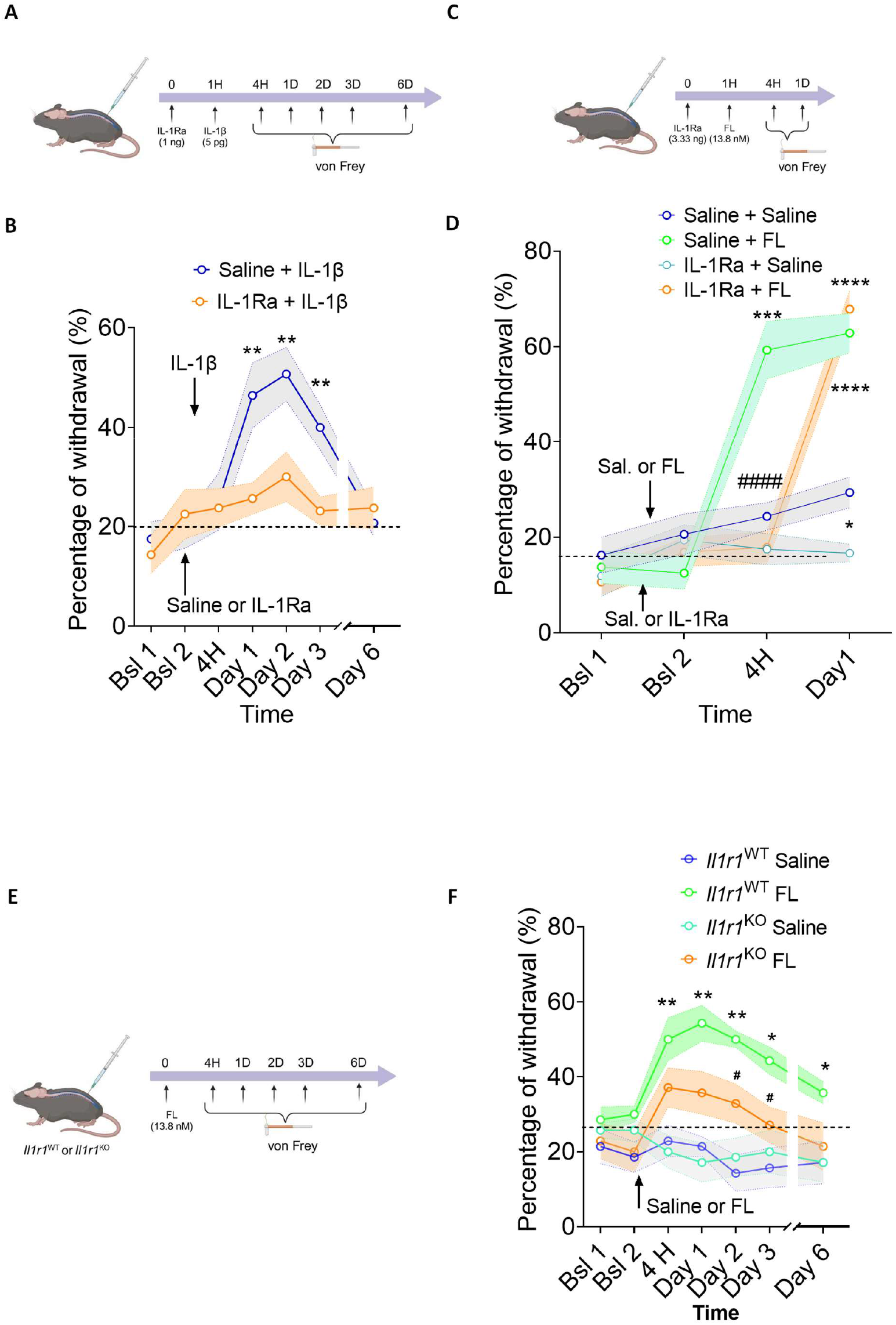
Effects of IL-1R signalling blockade on FL-induced mechanical pain hypersensitivity in mice. **A, C, E)** Schematic timelines representation of the experimental protocol on mice. **B)** Mechanical pain sensitivity, assessed as the percentage of withdrawal in response to von Frey filament application 4 hours to 6 Days after a double offbeat (20 minutes) intrathecal injection of Saline + IL-1β (5 pg/10 µl) or IL-1Ra (1 ng/10 µl) + IL-1β in Wild Type mice. **D)** Mechanical pain sensitivity, assessed as the percentage of withdrawal in response to Von Frey filament application 4 hours to 1 Day after a double offbeat (20 minutes) intrathecal injection of Saline + Saline or Saline + FL (13.8 nM) or IL-1Ra (3.33 ng/10 µl) + Saline or IL-1Ra (3.33 ng/10 µl) + FL (13.8 nM) in Wild Type mice. Number of animals in per group n = 10, Statistical analysis: two-way ANOVA followed by Dunnett test (A) or Bonferroni test (B), * p value < 0.05; ** p value < 0.01; *** p value< 0.001, **** p value< 0.0001 (Vs saline and #### p value < 0,001 (Saline-FL Vs IL-1Ra-FL). **F)** Mechanical pain sensitivity, assessed as the percentage of withdrawal in response to Von Frey filament 4 hours to 6 Days after FL or Saline injection in WT or *IL-1R1*^*KO*^ mice. Number of animals per group n = 7, Statistical analysis: two-way ANOVA followed by Dunnett multiple comparisons test * P < 0.05 and ** P < 0.01 (WT FL Vs WT saline group) and # P < 0,05 (*IL-1R1*^*KO*^ FL group versus WT FL group). Bsl Baseline before injection.

Further results showing mechanical pain hypersensitivity triggered by IT FL (13.8 nM/10µl) injection in the presence or absence of IL-1Ra (3.33 ng/10µl) are presented in **Figure 3C**. Paw withdrawal thresholds were evaluated over a 24-hour period using the Von Frey test. As observed in **Figure 3D**, IT injection of FL markedly increased the percentage of paw withdrawal compared to the Saline control group confirming FL-induced hyperalgesia as previously described (Rivat et al., 2018). Moreover, IL-1Ra (3.33 ng/10µl) injected one hour prior to FL injection significantly delayed the onset of FL-induced mechanical pain hypersensitivity. In the group treated with both IL-1Ra and FL the percentage of withdrawal responses induced by FL was near baseline level at 4H but similar to the control (Saline-FL group) at 24H **(Figure 3D)**. To further validate these findings, we investigated mechanical pain hypersensitivity triggered by IT FL (13.8 nM/10µl) injection in *Il1r1*^*KO*^ mice. Paw withdrawal thresholds were evaluated over a 6-day period using the Von Frey test **Figure 3E**. As shown in **Figure 3F**, FL injection induced a significant increase in mechanical pain in wild-type (*Il1r1*^WT^) mice compared to *Il1r1*^WT^ saline controls (**p < 0.05, p < 0.01**). In contrast, FL injection in *Il1r1*^*KO*^ had no significant effect compared to *Il1r1*^*KO*^ saline controls over the 6-day period. This effect was significantly lower than that induced by FL in *Il1r1*^WT^ FL-injected mice from day 2 post-injection (**p < 0.05**). Collectively our data suggest that the early pronociceptive effect of FL is dependent on IL-1β activity *in vivo*.

### 3.4 IL-1β induces mechanical pain hypersensitivity in *Flt3*^KO^ mice

**Figure 4A** illustrates a time-course experiment showing the mechanical pain hypersensitivity induced by IT injection of IL-1β (10 pg/10µl), in *Flt3* knockout (*Flt3*^KO^) mice, in comparison to wild-type (*Flt3*^WT^) mice. Paw withdrawal thresholds were measured using the von Frey test over a 6-day period. The results indicate that IL-1β induces similar mechanical pain hypersensitivity responses peaking at 4 hours post-injection, in both strains of mice (**Figure 4B**). These data strongly suggest that the pronociceptive effect of IL-1β is likely mediated through mechanisms that do not depend on FLT3 receptor presence or activation.

**Figure 4:**
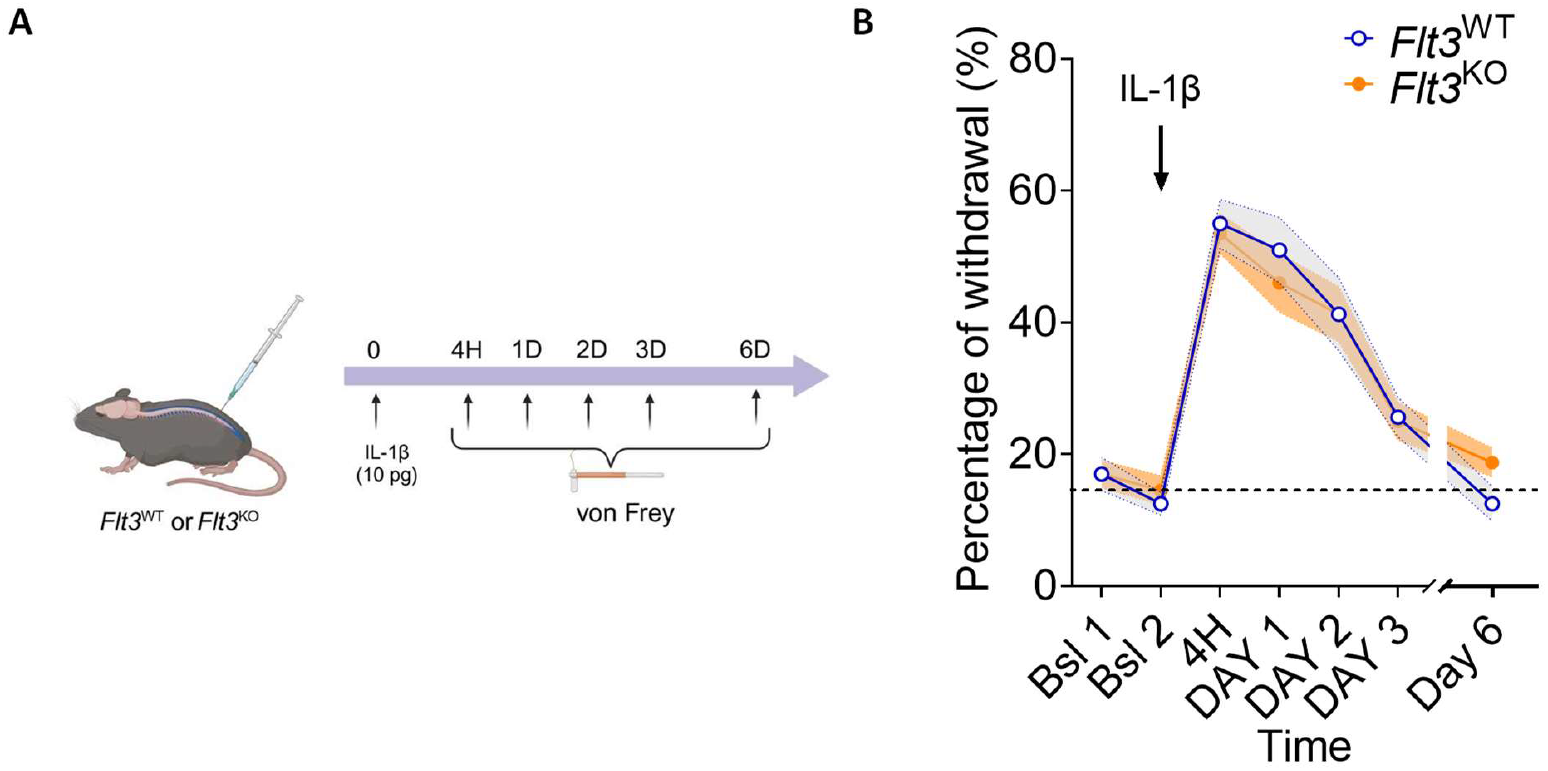
IL-1β-induced mechanical pain hypersensitivity in *Flt3* Wild Type or *Flt3* KO mice. **A)** Schematic timeline representation of the experimental protocol on mice. **B)** Mechanical pain sensitivity, assessed as the percentage of withdrawal to in response to Von Frey filament application, 5 to 144 hours after a single intrathecal of 10 pg/10 µl of IL-1β injection in *Flt3* Wild Type (*Flt3*^*WT*^) versus Flt3 Knock Out (*Flt3*^*KO*^) mice. Bsl Baseline before injection. Number of animals per group n = 10, Statistical analysis: two-way ANOVA, p value > 0.05 (NS).

## Discussion

Neuro-immune crosstalk, involving cytokines and neural pathways, significantly impacts pain perception and sensory processing, especially in inflammatory conditions (Marchand et al., 2005). In this study, we explored the potential interaction between FLT3 receptor activation by its ligand FL and the IL-1β/IL-1R signalling pathway; two processes known to play critical roles in neuropathic pain sensitization (Ren and Torres, 2009; Rivat et al., 2018). Our combined *in vitro* and *in vivo* findings provide compelling evidence that the early pronociceptive effects of FL are dependent on IL-1β signalling, thereby expanding our current understanding of cytokine interactions in pain mechanisms.

Cytokines, such as IL-1β, IL-6, and TNF-α, are known to engage in complex interactions and regulate each other, especially within the framework of immune responses and inflammation (Zhang and An, 2007). While FL is primarily known for its critical role in haematopoiesis and leukemogenesis through its interaction with the FLT3 receptor (Gilliland and Griffin, 2002; Kiyoi et al., 2020), such an interplay has yet to be explored in the context of neuroinflammation-related alterations in pain sensitivity. We first demonstrate that intrathecal administration of recombinant FL, significantly increased total IL-1β protein levels in both DRG and DSC as early as 4 hours post-injection. This early effect of FL on IL-1β expression appears to be unidirectional, as IL-1β did not significantly alter FL protein levels at this time point. Several cytokines are known to induce their own production as well as that of other cytokines. Since IL-1β can be produced by various cell types (Ren and Torres 2009 for review) it is reasonable to hypothesize that FL signalling may indirectly increase IL-1β from glial or immune cells in the DRG, or directly through neuronal pathways. Indeed, IL-1β and IL-1R expression has been described in rat and mouse DRG, in neuronal and non-neuronal cells, with likely co-localization within the same neurons indicating either an autocrine or a paracrine mechanism of interaction (Copray, 2001; Li et al., 2005; Fujita et al., 2024).

The interaction between FL and IL-1β signalling is also supported by our *in vitro* calcium imaging studies. Our data demonstrate that both FL and IL-1β enhanced TRPV1-mediated calcium influx in DRG neurons, a key effect involved in pain sensitization (Caterina et al., 2000; Davis et al., 2000; Basbaum et al., 2009). Furthermore, our findings show that pre-treatment with the IL-1R antagonist, IL-1Ra either completely blocked or significantly reduced the potentiated TRPV1 response induced by FL or IL-1β. While confirming the specificity of IL-1Ra, these results suggest that FL effect on TRPV1 potentiation is at least partially mediated through IL-1β/IL-1RI signalling pathways, raising questions about the underlying mechanisms.

Our calcium imaging studies were conducted using *in vitro* DRG cell cultures containing sensory neurons satellite glial and immune cells. Notably, IL-1R1 is known to be expressed in TRPV1^+^ DRG neurons in mice (Mailhot et al., 2020). Our current study further supports our previous observation that the FLT3 receptor regulates the activity of TRPV1+ neurons (Rivat et al., 2018). Moreover, our new findings suggest that FL-induced activation of the FLT3 receptor in TRPV1+ sensory neurons may initiate IL-1β/IL-1R1 signalling, with IL-1β released in its bioactive form following FLT3 activation. The potentiation of heat-activated ionic currents in nociceptors *via* an IL-1 R1-mediated mechanism has been reported in rats and could potentially represent a similar underlying mechanism in our model (Obreja et al., 2002).

The interaction between IL-1β and FL is further supported by our *in vivo* experiments. Our data indicate that pre-treatment with IL-1Ra, administered one hour before FL in naïve mice, results in several hours delay in FL-induced mechanical pain hypersensitivity. This finding suggests that IL-1β signalling is involved in FL-induced pain responses *in vivo*. Although this delay might be related to the half-life of the antagonist IL-1Ra, it could also indicate a critical window for IL-1β activity following FLT3 activation. Supporting this latter hypothesis, administration of IL-1Ra 20 hours post-FL injection had no effect on FL-induced pain hypersensitivity (data not shown). Moreover, FL-induced mechanical pain hypersensitivity in mice was evaluated in *Il1r1*^*KO*^. The lack of a significant pain effect of FL in this condition strengthens our conclusions that IL-1β signalling is involved in FL-induced pain responses *in vivo*. The effects observed in *Flt3*^KO^ mice further suggest that the IL-1β acute pronociceptive effects seems to be mechanistically independent of FLT3, offering insights into distinct yet interacting molecular mechanisms implicated in neuropathic pain.

Overall, our study extends on our knowledge on neuro-immune cross talks by demonstrating the functional link between FL/FLT3 and IL-1β/IL-1R signalling in nociceptive pathways. Notably, it highlights IL-1β critical role in mediating FL activation-induced mechanical pain hypersensitivity through IL-1R signalling in normal animals and open the possibility to test this hypothesis in pain syndrome where both cytokines are implicated.

## References

Arend, W. P. (1991). Interleukin 1 receptor antagonist. A new member of the interleukin 1 family. J. Clin. Invest. 88, 1445–1451. doi: 10.1172/JCI115453

Basbaum, A. I., Bautista, D. M., Scherrer, G., and Julius, D. (2009). Cellular and Molecular Mechanisms of Pain. Cell 139, 267–284. doi: 10.1016/j.cell.2009.09.028

Binshtok, A. M., Wang, H., Zimmermann, K., Amaya, F., Vardeh, D., Shi, L., et al. (2008). Nociceptors Are Interleukin-1β Sensors. J. Neurosci. 28, 14062–14073. doi: 10.1523/JNEUROSCI.3795-08.2008

Caterina, M. J., Leffler, A., Malmberg, A. B., Martin, W. J., Trafton, J., Petersen-Zeitz, K. R., et al. (2000). Impaired Nociception and Pain Sensation in Mice Lacking the Capsaicin Receptor. Science 288, 306–313. doi: 10.1126/science.288.5464.306

Chu, C., Artis, D., and Chiu, I. M. (2020). Neuro-immune Interactions in the Tissues. Immunity 52, 464–474. doi: 10.1016/j.immuni.2020.02.017

Copray, J. (2001). Expression of interleukin-1 beta in rat dorsal root ganglia. Journal of Neuroimmunology 118, 203–211. doi: 10.1016/S0165-5728(01)00324-1

Dantzer, R. (2018). Neuroimmune Interactions: From the Brain to the Immune System and Vice Versa. Physiological Reviews 98, 477–504. doi: 10.1152/physrev.00039.2016

Davis, J. B., Gray, J., Gunthorpe, M. J., Hatcher, J. P., Davey, P. T., Overend, P., et al. (2000). Vanilloid receptor-1 is essential for inflammatory thermal hyperalgesia. Nature 405, 183–187. doi: 10.1038/35012076

Elzière, L., Sar, C., Ventéo, S., Bourane, S., Puech, S., Sonrier, C., et al. (2014). CaMKK-CaMK1a, a New Post-Traumatic Signalling Pathway Induced in Mouse Somatosensory Neurons. PLoS ONE 9, e97736. doi: 10.1371/journal.pone.0097736

Fujita, D., Matsuoka, Y., Yamakita, S., Horii, Y., Ishikawa, D., Kushimoto, K., et al. (2024). Rapid cleavage of IL-1β in DRG neurons produces tissue injury-induced pain hypersensitivity. Mol Pain 20, 17448069241285357. doi: 10.1177/17448069241285357

Gilliland, D. G., and Griffin, J. D. (2002). The roles of FLT3 in hematopoiesis and leukemia. Blood 100, 1532–1542. doi: 10.1182/blood-2002-02-0492

Glaccum, M. B., Stocking, K. L., Charrier, K., Smith, J. L., Willis, C. R., Maliszewski, C., et al. (1997). Phenotypic and functional characterization of mice that lack the type I receptor for IL-1. J Immunol 159, 3364–3371.

Grace, P. M., Hutchinson, M. R., Manavis, J., Somogyi, A. A., and Rolan, P. E. (2010). A novel animal model of graded neuropathic pain: Utility to investigate mechanisms of population heterogeneity. Journal of Neuroscience Methods 193, 47–53. doi: 10.1016/j.jneumeth.2010.08.025

Kiyoi, H., Kawashima, N., and Ishikawa, Y. (2020). FLT3 mutations in acute myeloid leukemia: Therapeutic paradigm beyond inhibitor development. Cancer Sci 111, 312– 322. doi: 10.1111/cas.14274

Li, M., Shi, J., Tang, J., Chen, D., Ai, B., Chen, J., et al. (2005). Effects of complete Freund’s adjuvant on immunohistochemical distribution of IL-1beta and IL-1R I in neurons and glia cells of dorsal root ganglion1. Acta Pharmacologica Sinica 26, 192–198. doi: 10.1111/j.1745-7254.2005.00522.x

Mailhot, B., Christin, M., Tessandier, N., Sotoudeh, C., Bretheau, F., Turmel, R., et al. (2020). Neuronal interleukin-1 receptors mediate pain in chronic inflammatory diseases. Journal of Experimental Medicine 217, e20191430. doi: 10.1084/jem.20191430

Marchand, F., Perretti, M., and McMahon, S. B. (2005). Role of the Immune system in chronic pain. Nat Rev Neurosci 6, 521–532. doi: 10.1038/nrn1700

McMahan, C. J., Slack, J. L., Mosley, B., Cosman, D., Lupton, S. D., Brunton, L. L., et al. (1991). A novel IL-1 receptor, cloned from B cells by mammalian expression, is expressed in many cell types. The EMBO Journal 10, 2821–2832. doi: 10.1002/j.1460-2075.1991.tb07831.x

Moalem, G., and Tracey, D. J. (2006). Immune and inflammatory mechanisms in neuropathic pain. Brain Research Reviews 51, 240–264. doi: 10.1016/j.brainresrev.2005.11.004

Obreja, O., Rathee, P. K., Lips, K. S., Distler, C., and Kress, M. (2002). IL-1J potentiates heat-activated currents in rat sensory neurons: involvement of IL-1RI, tyrosine kinase, and protein kinase C. The FASEB Journal 16, 1497–1503. doi: 10.1096/fj.02-0101com

Orlando, S., Sironi, M., Bianchi, G., Drummond, A. H., Boraschi, D., Yabes, D., et al. (1997). Role of Metalloproteases in the Release of the IL-1 type II Decoy Receptor. Journal of Biological Chemistry 272, 31764–31769. doi: 10.1074/jbc.272.50.31764

Ren, K., and Torres, R. (2009). Role of interleukin-1β during pain and inflammation. Brain Research Reviews 60, 57–64. doi: 10.1016/j.brainresrev.2008.12.020

Rivat, C., Sar, C., Mechaly, I., Leyris, J.-P., Diouloufet, L., Sonrier, C., et al. (2018). Inhibition of neuronal FLT3 receptor tyrosine kinase alleviates peripheral neuropathic pain in mice. Nat Commun 9, 1042. doi: 10.1038/s41467-018-03496-2

Zhang, J.-M., and An, J. (2007). Cytokines, Inflammation, and Pain. International Anesthesiology Clinics 45, 27–37. doi: 10.1097/AIA.0b013e318034194e

